# Monothiol glutaredoxin DaGrxB couples iron homeostasis with redox balance to orchestrate development and virulence in the peach shoot blight fungus *Diaporthe amygdali*

**DOI:** 10.64898/2026.09.28.755239

**Authors:** Lina Yang, Xun Liu, Lili Xiong, Yue Zhang, Yaru Jing, Hengsong Shi, Ziyi Pan, Shiyuan Cao, Lianwei Li, You Liang, Zhaolin Ji

## Abstract

Peach shoot blight (PSB), caused by *Diaporthe amygdali*, is one of the most devastating fungal diseases affecting peach production worldwide. Glutaredoxins (Grxs) are ubiquitous redox-regulatory proteins involved in antioxidant defense and iron metabolism, yet their functions in phytopathogenic fungi remain largely unknown. Here, we identified four Grxs (DaGrxA-DaGrxD) in *D. amygdali*. Among them, DaGrxB, a monothiol glutaredoxin, is the sole member essential for vegetative growth, asexual reproduction and virulence. DaGrxB interacts with iron-regulatory transcription factors DaHapX and DaSreA and modulates their activities via iron-dependent nucleocytoplasmic shuttling, coordinating iron acquisition and storage gene expression. During infection, *D. amygdali* encounters a biphasic iron environment: an inherently iron-limited host environment at early stages (1-2 dpi) followed by host-mediated iron excess (3-4 dpi), and DaGrxB is essential for navigating this transition. Consistently, deletion of either *DaHAPX* or *DaSREA* significantly reduced virulence, confirming that iron homeostasis is a critical determinant of *D. amygdali* virulence. Notably, deletion of *DaGRXB* disrupts Fe-S cluster biosynthesis, causing mitochondrial Fe²⁺ accumulation, impaired aconitase activity, and reduced ATP production, which triggers constitutive activation of the GSH-dependent antioxidant system, with elevated GSH/GSSG ratios, increased GR and GST activities, and reduced H₂O₂ levels. Exogenous GSH and H₂O₂ depletion both reduced virulence, suggesting that disturbed redox balance directly contributes to attenuated virulence. Additionally, the Trx domain assists the Grx domain in nucleocytoplasmic translocation through intramolecular interactions. Finally, LDH-delivered dsRNA targeting *DaGRXB* effectively controls PSB. Collectively, these findings reveal how a monothiol glutaredoxin integrates iron homeostasis, redox balance and virulence in a plant pathogen, providing a foundation for RNAi-based disease management.

**Author summary:** Peach shoot blight is a devastating disease that causes substantial yield losses in peach production worldwide. The causal fungus, *Diaporthe amygdali*, also infects other economically important fruit crops. During infection, plants use iron as a weapon: either by starving the pathogen of iron or by releasing toxic levels of it. Glutaredoxins (Grxs) are ubiquitous redox-regulatory proteins involved in antioxidant defense and iron metabolism, and they are emerging as key players in this process. We studied how this fungus fights back. We found that it carries four glutaredoxin (Grx) proteins, but only DaGrxB is essential for its development and virulence. DaGrxB acts as a molecular switch that helps the fungus sense and respond to the changing iron levels inside the host. It does this by moving between the nucleus and cytoplasm, where it works together with two iron-regulatory proteins DaHapX and DaSreA to control iron uptake and storage genes expression. When we deleted *DaGrxB*, the fungus could no longer manage iron properly, leading to disrupted energy production and an overactive antioxidant system. Importantly, we also developed a spray that uses RNA interference to target *DaGRXB*, which effectively controlled the disease. Our findings reveal how a single protein helps a fungal pathogen survive the host’s iron-based defenses, and offer a promising strategy for disease management.

## Introduction

Peach shoot blight (PSB), caused by *Diaporthe amygdali*, is one of the most devastating fungal diseases affecting peach production worldwide, causing yield losses of 20-50% and complete crop failure in severe cases(Lalancette, 2003; Froelich and Schnabel, 2019; Beluzán et al., 2021; Brugneti et al., 2025). The pathogen also infects other economically important fruit crops, including pear, plum, almond, and apple(Francia et al., 2025). As an opportunistic pathogen, *D. amygdali* enters host tissues through wounds under conditions of high temperature and humidity, infecting peach buds, twigs, and fruits, but not leaves. Despite its economic importance, the molecular mechanisms underlying *D. amygdali* pathogenesis remain poorly understood. Our previous work characterized nine MAPK cascade components and velvet family proteins in this pathogen(Yang et al., 2025; Shi et al., 2026), providing a foundation for further functional studies.

Reactive oxygen species (ROS) cause oxidative damage to biological macromolecules, and cells have evolved redox-regulatory proteins to maintain homeostasis. Glutaredoxins (Grxs) and thioredoxins (Trxs) are two major families of oxidoreductases central to the reduction of oxidized proteins, with Grxs exhibiting broader substrate versatility(Grant, 2001; Herrero et al., 2010). Ubiquitous in nature, Grxs are involved in antioxidant defense and iron metabolism(Fernandes and Holmgren, 2004; Berndt et al., 2021). Originally identified as glutathione (GSH)-dependent electron donors for ribonucleotide reductase(Holmgren, 1976), Grxs are now recognized as central regulators of cellular redox homeostasis, operating through a conserved GSH/glutathione reductase(GR)/NADPH system(Cao et al., 2020; Ogata et al., 2021).Their hallmark function is to catalyze reversible protein glutathionylation, a post-translational modification that dynamically modulates protein function(Guo et al., 2021). Based on active-site motif, Grxs are divided into two major groups: Class I (CXXC/S motif) comprises oxidoreductases, while Class II (CGFS motif) comprises monothiol Grxs specialized in iron homeostasis and iron-sulfur (Fe-S) cluster biosynthesis(Herrero et al., 2006; Mühlenhoff et al., 2010; Li and Outten, 2012). Although well characterized in model organisms, the roles of Grxs in plant pathogenic fungi are only beginning to emerge. For instance, monothiol glutaredoxin Grx4 is essential for virulence on *Ustilago maydis*(McCotter et al., 2023), AaGrxD contributes to virulence in *Alternaria alternata*(Lu et al., 2026), and Grx3/4 supports full virulence in human fungal pathogen *Candida albicans*(Snyder et al., 2020) and *Cryptococcus neoformans*(Attarian et al., 2018). Nevertheless, the functions of Grx proteins in phytopathogenic fungi remain largely unknown.

Iron is an essential micro-nutrient and co-factor for numerous enzymes involved in energy metabolism, biosynthesis, replication, and cellular redox status(Frey and Reed, 2012; Crielaard et al., 2017). In plant-pathogen interactions, iron availability constitutes a critical battleground. Plants employ two opposing strategies to restrict pathogens: nutritional immunity, which limits iron availability to inhibit growth, and Fenton-based defense, which locally elevates iron to generate toxic hydroxyl radicals that directly damage pathogen cells(Kieu et al., 2012; Ye et al., 2014).To counteract these host-imposed stresses, plant pathogens have evolved sophisticated regulatory networks to sense and respond to fluctuating iron levels.

In filamentous fungi, iron homeostasis is primarily governed by two transcription factors: the bZIP protein HapX and the GATA-type repressor SreA. These regulators coordinate genes involved in iron acquisition (siderphore-mediated SIA and reductive RIA pathways), utilization (TCA cycle, respiration, heme metabolism, and Fe-S cluster biosynthesis), and storage (vacuolar sequestration) (Haas, 2012; Wiemann et al., 2014). Under iron starvation, HapX activates iron acquisition pathway while repressing iron-consuming pathways and *SREA* expression; under iron-replete conditions, HapX activates iron storage and utilization, whereas SreA represses iron acquisition to prevent overload(Cowen et al., 2010; Haas, 2012; Wiemann et al., 2014). The contributions of HapX and SreA to virulence are functionally asymmetric and species-specific. Deletion of *HAPX* attenuates virulence in *Aspergillus fumigatus*, *Fusarium oxysporum*, *Magnoporthe oryzae* and *Valsa dahliae*, whereas deletion of *SREA* does not affect virulence in *A. fumigatus* or *A. alternata*(Cowen et al., 2010; López-Berges et al., 2012; Kong et al., 2014; Wang et al., 2018; Chung et al., 2020). Notably, in *Fusarium graminearum*, inactivation of either regulator reduces virulence(Sun et al., 2023), suggesting context-dependent importance.

Despite these established roles, the upstream signaling regulators that coordinating HapX and SreA in plant pathogenic fungi remain poorly understood. Recent evidence implicated FgHapX in activating iron storage genes via histone H2B deubiquitination, while FgSreA represses iron acquisition genes via H2A.Z deposition and H3K27 trimethylation under iron excess in *F. graminearum*(Sun et al., 2023). Moreover, the monothiol glutaredoxin FgGrx4 controls the transcriptional activity of both regulators by modulating their enrichment at target genes and recruiting epigenetic regulators(Sun et al., 2023). However, whether and how Grxs regulate HapX/SreA-mediated iron homeostasis in phytopathogenic fungi remains elusive.

In this study, we identified four Grxs in *D. amygdali* and demonstrated that DaGrxB, a monothiol glutaredoxin, is the only member required for vegetative growth, asexual reproduction, and virulence. DaGrxB physically interacted with DaHapX and DaSreA and modulates their activities through iron-dependent nucleocytoplasmic shuttling, coordinating iron acquisition and storage to adapt to dynamic iron environments during infection. Deletion of *DaGRXB* impaired Fe-S cluster biosynthesis and activation of the GSH-dependent antioxidant system. We further revealed that the Trx and Grx domains of DaGrxB jointly regulated its biological function. Moreover, nanomaterial-delivered dsRNA targeting *DaGRXB* effectively controlled PSB. Taken together, our findings elucidate a novel regulatory mechanism of monothiol glutaredoxin in plant pathogenic fungi and provide a promising RNA interference (RNAi)-based strategy for disease management.

## Materials and Methods

### Strains and culture conditions

*D. amygdali* strain ZN32 (Yang et al., 2022) was used as the wild-type control. All strains were cultured on PDA, CM, OM, or AC media at 25°C in the dark. Transformants were selected on TB3 medium containing 300 μg/mL hygromycin B or 200 μg/mL zeocin. Stress sensitivity was assessed on CM medium supplemented with iron or oxidative stressors for 4 d. Peach cultivar ‘Liutiao Baifeng’ was used for inoculation.

### Generation of gene deletion mutants and complemented strains

Gene deletion mutants were generated via PEG-mediated protoplast transformation (Yang et al., 2025; Shi et al., 2026). The ORF of each target gene was replaced with an *HPH* cassette using a split-marker approach. Protoplasts of ZN32 were transformed with two overlapping PCR products and the corresponding pCas9-gRNA vector. Hygromycin B-resistant transformants were confirmed by PCR. For complementation, ORFs were cloned into *Xho*I-linearized pYF11 vectors under the RP27 promoter, with DaGrxA-DaGrxD fused upstream of GFP and DaHapX/DaSreA fused downstream of GFP. Plasmids were transformed into corresponding deletion mutants and selected on zeocin. Primers are listed in Table S1.

### Virulence assay

One-year-old peach twigs were surface-sterilized with 75% ethanol for 30 s, rinsed, air-dried, and cut into 7-cm segments with three equidistant wounds. Hyphal blocks (9 mm × 9 mm) were placed onto wounds and incubated at 28°C with 90% relative humidity (RH) under a 12-h light/12-h dark photoperiod for 7 d. Fruits were inoculated similarly and assessed at 5 dpi. Three independent replicates were performed.

### Co-IP assay

Total proteins were extracted from ZN32/DaGrxB-GFP and ZN32/GFP cultured in liquid CM at 28°C for 36 h using NP40 lysis buffer. Extracts were incubated with GFP-Trap beads overnight at 4°C. After washing, bound proteins were eluted with 0.2 M glycine (pH 2.5), neutralized with 1 M Tris-base, and analyzed by MS.

### Measurement of intracellular iron content

Mycelia were cultured in liquid CM at 28°C for 36 h. Mycelium (0.5 g) was digested with nitric perchloric acid until clear. Iron content was determined by atomic absorption spectrophotometry (AAS) at 248.3 nm. Content was calculated as: X (mg/kg) = (ρ – ρ₀) × V × F / M, where ρ and ρ₀ are sample and blank concentrations, V is final volume, M is sample mass, and F is the dilution factor. Three independent replicates were performed.

### qRT-PCR assay

For gene expression analysis, strains were cultured in liquid CM at 28°C for 36 h. For in planta assays, 10 μL of conidia (1 × 10⁶ spores/mL) from ZN32 were inoculated onto wounded peach fruits, and tissues were harvested at indicated times. Total RNA was extracted (PureLink RNA Mini Kit; Invitrogen) and reverse transcribed (HiScript II; Vazyme). Transcript levels were quantified with *DaACTIN* as internal control. Three independent replicates were performed. Primers are listed in Table S1.

### Mitochondrial Fe²⁺ staining

Mycelia were cultured in liquid CM at 28°C for 18 h. Pellets were washed, blotted dry, and stained with 30 μL Mito-FerroGreen2 (1000×; Beyotime) for 30 min in the dark. Staining was observed under confocal microscope (Ex/Em = 515/545 nm). At least 50 images were captured per strain, and the experiment was repeated independently three times.

### ACO activity measurement

ACO activity was measured using a kit (BC4485; Solarbio). For total ACO, mycelial powder (0.1 g) was homogenized with 1 mL Buffer 1 and 10 μL Buffer 3 on ice, sonicated (300 W, 3 s on/9 s off, 15 cycles), and centrifuged at 11,000 g for 15 min at 4°C. For fractionation, mycelial powder (0.2 g) was homogenized and centrifuged at 600 g for 5 min; the supernatant was further centrifuged at 11,000 g for 15 min as the cytoplasmic fraction, while the pellet was resuspended in 400 μL Buffer 2 with 4 μL Buffer 3, sonicated, and centrifuged at 5,000 g for 2 min as the mitochondrial fraction. Activity was measured by mixing 20 μL sample with 180 μL pre-warmed Buffer 4 (25°C, 15 min). Absorbance at 240 nm was recorded at 10 s (A₁) and 5 min 10 s (A₂). Activity (U/g) = 561.11 × (A₂ − A₁) ÷ W (total and cytoplasmic) or 244.44 × ΔA ÷ W (mitochondrial), where W is sample mass (g). Three independent replicates were performed.

### ATP content measurement

ATP was measured using a kit (S0026; Beyotime). Mycelial powder (0.1 g) was lysed with 100 μL lysis buffer and centrifuged at 12,000 g for 5 min at 4°C. In a well, 100 μL lysis buffer was incubated for 5 min, followed by 20 μL supernatant. Luminescence (RLU) was measured after 2 s. A standard curve was generated using serial dilutions of ATP standard. ATP content was normalized to protein concentration (BCA assay; Beyotime) and expressed as nmol/mg protein. Three independent replicates were performed.

### GSH and GSSG content measurement

Mycelia were cultured in liquid CM at 28°C for 36 h, harvested, and ground in liquid nitrogen. Powder (0.1 g) was homogenized with 250 μL 5% metaphosphoric acid (MPA) buffer on ice. After centrifugation (8,000 g, 10 min, 4°C), supernatant was collected. For GSH, 20 μL supernatant was mixed with 140 μL phosphate buffer (pH 7.0) and 40 μL 4 mM 5,5′-dithiobis (2-nitrobenzoic acid) (DTNB); absorbance at 412 nm (A₁) was read after 2 min against a blank (A₀); ΔA = A₁ − A₀. For GSSG, 20 μL supernatant was incubated with 400 μL 2-vinylpyridine at 37°C for 30 min, then mixed with 140 μL phosphate buffer (pH 7.5), 20 μL NADPH, 20 μL DTNB, and 2 μL GR; absorbance was recorded at 412 nm at 30 s (A₁) and 5 min 30 s (A₂); ΔA₁ = A₂ − A₁. GSH and GSSG contents were calculated using standard curves with the formula: GSH or GSSG (μg/g) = C × V × F / M, where C is concentration from standard curves (μg/mL), V is buffer volume (0.25 mL), M is sample mass (g), and F is dilution factor. Three independent replicates were performed.

### GR and GST activity assays

GR activity was measured using a kit (GR-2-W; Comin). Powder (0.1 g) was homogenized with 1 mL Buffer 1 on ice and centrifuged at 8,000 g for 15 min at 4°C. Supernatant (100 μL) was mixed with 750 μL Buffer 1, 100 μL Buffer 2, and 50 μL Buffer 3. Absorbance at 340 nm was recorded at 0 s (A₁) and 180 s (A₂). GR activity (nmol/min/g) = 536 × (A₁ − A₂) ÷ W.

GST activity was measured using a kit (GST-2-W; Comin). Powder (0.1 g) was homogenized with 1 mL Buffer 1 on ice and centrifuged at 8,000 g for 10 min at 4°C. Supernatant (100 μL) was mixed with 900 μL Buffer 2 and 100 μL Buffer 3. Absorbance at 340 nm was recorded at 10 s (A₁) and 310 s (A₂). GST activity (nmol/min/g) = 230 × (A₂ − A₁) ÷ W. Three independent replicates were performed for each assay.

### H₂O₂ content measurement

Powder (0.1 g) was homogenized with 500 μL 1× PBS and centrifuged at 8,000 g for 10 min at 4°C. Supernatant (250 μL) was mixed with 50 μL concentrated ammonia and 25 μL titanium sulfate solution (50 mg/mL), then centrifuged at 8,000 g for 10 min at 25°C. The precipitate was dissolved in 250 μL 2 M H₂SO₄. After 5 min, absorbance was measured at 415 nm (A₁) against an acetone blank (A₀); ΔA = A₁ − A₀. A standard curve was generated using serial dilutions of H₂O₂ standard. H₂O₂ content (μmol/g) = C × V × F / M, where C is from the standard curve (μmol/mL), V is extraction volume (0.5 mL), M is sample mass (g), and F is dilution factor. Three independent replicates were performed.

### Spray-induced gene silencing (SIGS) assay

dsRNA targeting *DaGRXB* (20-420 bp) was synthesized *in vitro* (HighYield T7 RNA Transcription kit, SJ002; Belongbio) and purified (PU002; Belongbio). dsRNA^GFP^ was used as negative control. Nanomaterial layered double hydroxide (LDH) was provided by Liang You. dsRNA (50 ng/μL) was mixed with LDH at a 1:4 (w/w) ratio. Wounded twigs (2 × 4 mm) were treated with 10 μL dsRNA (50 ng/μL), dsRNA-LDH, or sterile water for 2 h, then inoculated with hyphal blocks (9 mm × 9 mm) of ZN32 and incubated at 28°C with 90% RH (12 h light/dark) for 7 d. Fungal biomass was determined by DNA-based qRT-PCR targeting *DaACTIN* and normalized to *PpTEF2*. *DaGRXB* expression was assessed by qRT-PCR and normalized to *DaACTIN*. Three independent replicates were performed.

## Results

### Identification of Grxs from *D. amygdali*

To identify Grx-encoding genes in *D. amygdali*, we performed BLASTp searches against the genome database using all eight *Saccharomyces cerevisiae* Grx proteins as queries. This identified four candidate genes, *GME11839_g*, *GME1975_g*, *GME7338_g*, and *GME4946_g*, encoding full-length Grx homologs. Phylogenetic analysis resolved these proteins into four clades, designated DaGrxA-DaGrxD. Multiple sequence alignment revealed conserved active-site motifs: CPYC (DaGrxA), CGFS (DaGrxB and DaGrxC), and CXXS (DaGrxD) (Fig S1A). Domain architecture analysis showed that all four proteins contain a single Grx domain; notably, DaGrxB possesses an additional N-terminal thioredoxin (Trx) domain, whereas DaGrxD features an N-terminal transmembrane domain (Fig S1B).

### Expression patterns of *DaGRX* genes

Transcript levels of the four *DaGRX* genes were quantified during developmental stages and in response to infection using qRT-PCR. *DaGRXA* expression remained unchanged across all tested conditions (Fig S2A). *DaGRXB* was strongly upregulated during conidiation (9.1-fold) and further induced during infection, peaking at 24 hpi (153.3-fold) (Fig S2B). *DaGRXC* showed no change during asexual development but was downregulated by ∼30% during infection (Fig S2C). *DaGRXD* was significantly induced during infection, peaking at 24 hpi (32.5-fold) (Fig S2D). These results suggest that DaGrxB functions in both asexual reproduction and host infection, whereas DaGrxD appears specifically involved in infection.

### DaGrxB is specifically required for vegetative growth and asexual reproduction

We generated single-gene deletion mutants (Δ*DagrxA*, Δ*DagrxB*, Δ*DagrxC*, and Δ*DagrxD*) via homologous recombination (Fig S3A, B) and complemented strains expressing GFP-tagged coding sequences under the RP27 promoter. On PDA, CM, OM, and AC media, Δ*DagrxB* mutants exhibited significantly reduced growth rates and thinner mycelium compared with the wild type ZN32, whereas other mutants showed no obvious differences (Fig 1A; Table 1). At 15 dpi on PDA, ZN32 and all mutants except Δ*DagrxB* produced abundant melanized pycnidia. Δ*DagrxB* mutants formed markedly fewer and smaller pycnidia, with counts at 15 and 30 dpi being ∼7% and ∼25% of wild-type levels. Moreover, while all other strains secreted copious α-conidia at 30 dpi, Δ*DagrxB* mutants failed to release conidia spontaneously; manual rupture revealed both α- and β-conidia (Fig 1B; Table 1), indicating that DaGrxB is required for pycnidium maturation and conidial discharge but not for conidiogenesis. These results demonstrate that DaGrxB, but not DaGrxA, DaGrxC, or DaGrxD, is essential for vegetative growth, pycnidium development, and conidial release.

**Fig 1.**
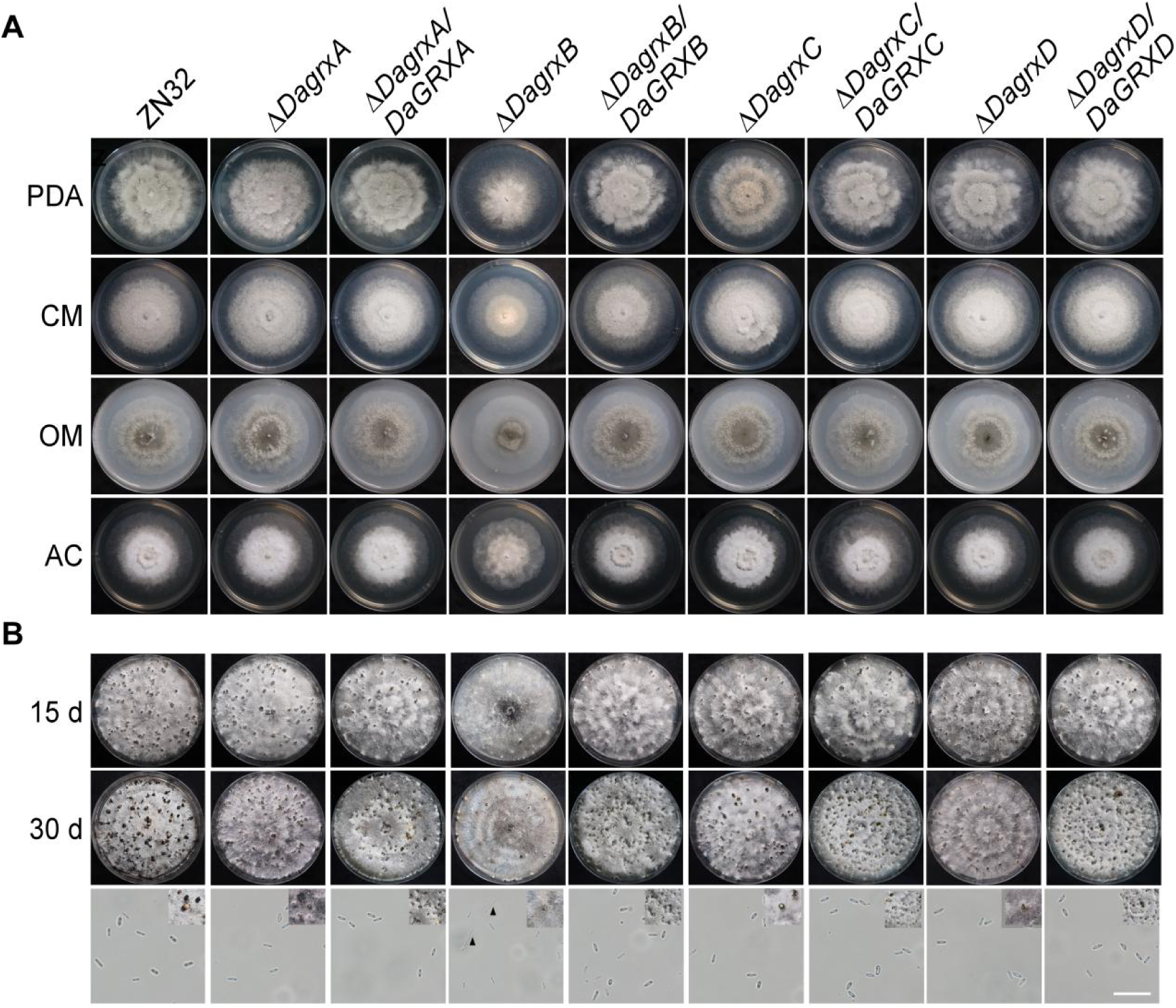
DaGrxB is specifically required for vegetative growth and asexual reproduction. (A) Colony morphology of ZN32, Δ*DagrxA*, Δ*DagrxB*, Δ*DagrxC*, Δ*DagrxD* mutants, and complemented strains on PDA, CM, OM, and AC medium at 25°C for 4 d. (B) Pycnidia formation on PDA at 15 and 30 d; conidial morphology (40×) of indicated strains, with black triangles indicating β-conidia. Bar = 20 μm.

**Table 1.** Phenotypic characterization of *DaGRXs* deletion mutants of *D. amygdali*.

| Strain | Colony diameters (cm)<sup>I</sup> |  |  |  | Pycnidia number <sup>II</sup> |  |
| --- | --- | --- | --- | --- | --- | --- |
|  | PDA | CM | OM | AC | 15 d | 30 d |
| ZN32 | 8.1 ± 0.1a | 8.0 ± 0.1a | 6.5 ± 0.1a | 6.6 ± 0.1a | 57 ± 2.5a | 76 ± 8.5a |
| $\Delta DagrxA$ | 8.0 ± 0.1a | 8.1 ± 0.3a | 6.4 ± 0.2a | 6.5 ± 0.1a | 50 ± 2.5a | 70 ± 7.6a |
| $\Delta DagrxA/DaGRXA$ | 8.1 ± 0.1a | 8.0 ± 0.1a | 6.5 ± 0.1a | 6.6 ± 0.1a | 49 ± 9.3a | 64 ± 9.6a |
| $\Delta DagrxB$ | 7.2 ± 0.1b | 6.1 ± 0.2b | 5.9 ± 0.1b | 5.7 ± 0.1b | 4 ± 5.1b | 19 ± 8.3b |
| $\Delta DagrxB/DaGRXB$ | 8.1 ± 0.1a | 8.1 ± 0.1a | 6.5 ± 0.1a | 6.5 ± 0.2a | 44 ± 8.5a | 60 ± 4.9a |
| $\Delta DagrxC$ | 8.0 ± 0.1a | 8.0 ± 0.1a | 6.5 ± 0.1a | 6.6 ± 0.1a | 46 ± 8.6a | 70 ± 2.5a |
| $\Delta DagrxC/DaGRXC$ | 8.1 ± 0.1a | 8.0 ± 0.1a | 6.5 ± 0.1a | 6.6 ± 0.1a | 49 ± 7.5a | 68 ± 7.0a |
| $\Delta DagrxD$ | 8.1 ± 0.1a | 8.0 ± 0.1a | 6.5 ± 0.1a | 6.6 ± 0.1a | 44 ± 4.7a | 58 ± 7.5a |
| $\Delta DagrxD/DaGRXD$ | 8.1 ± 0.1a | 8.0 ± 0.1a | 6.5 ± 0.1a | 6.7 ± 0.1a | 45 ± 5.3a | 61 ± 6.7a |

### DaGrxB is the sole Grx contributing to virulence

Peach twigs and fruits were inoculated with hyphal blocks of ZN32, each deletion mutant, and corresponding complemented strains. At 7 dpi, lesions caused by ZN32, Δ*DagrxA*, Δ*DagrxC*, and Δ*DagrxD* mutants, and all complemented strains averaged 1.9-2.0 cm on peach twigs. By contrast, Δ*DagrxB* mutants produced lesions of only 0.2 cm (Fig 2A, B), with fungal biomass reduced by ∼90% (Fig 2C). Similar results were obtained on peach fruits (Fig 2D-F). These findings establish that DaGrxB is indispensable for full virulence, whereas the other three Grxs are dispensable.

**Fig 2.**
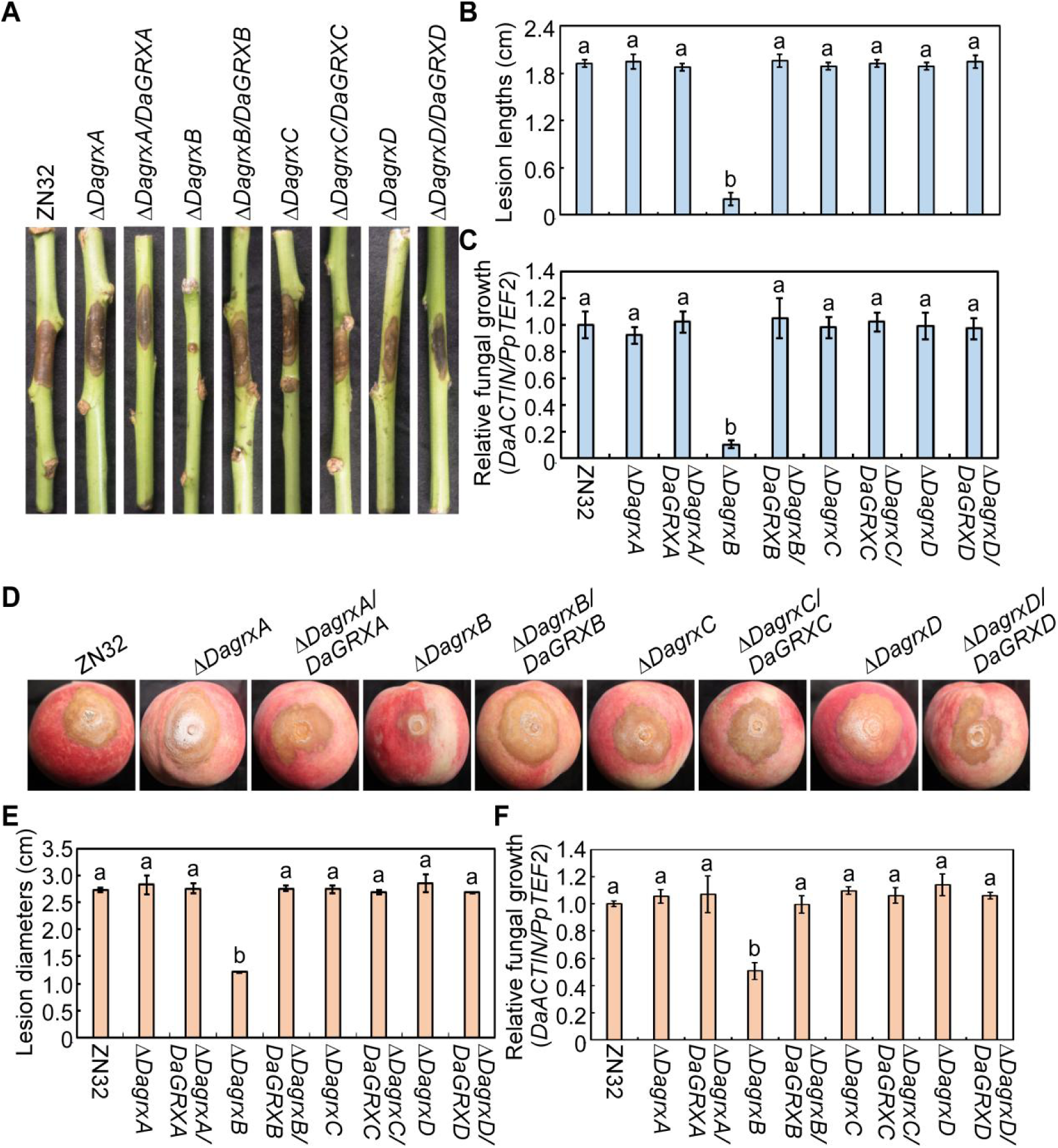
DaGrxB is the sole glutaredoxin required for *D. amygdali* virulence. (A) Disease symptoms on peach twigs at 7 dpi. (B) Lesion lengths. (C) Fungal biomass in twigs (qRT-PCR of *DaACTIN*, normalized to *PpTEF2*). (D) Disease symptoms on peach fruits at 5 dpi. (E) Lesion diameters. (F) Fungal biomass in fruits as in (C). Error bars, ±SD (n = 3). Different lowercase letters indicate significant differences (one-way ANOVA with Tukey’s HSD test, *P* < 0.05).

### Subcellular localization of DaGrx proteins

Confocal microscopy revealed that DaGrxA and DaGrxC localized exclusively to mitochondria, confirmed by colocalization with MitoTracker (Fig S4A-C). DaGrxB exhibited nucleocytoplasmic distribution in both hyphae and conidia (Fig S4A, B, D). DaGrxD displayed stage-specific localization: predominantly at septa in hyphae, but uniformly distributed in conidia (Fig S4A, B).

### DaGrxB interacts with DaHapX and DaSreA and is required for iron stress responses

Co-immunoprecipitation (Co-IP) assay coupled with mass spectrometry (MS) using GFP-tagged DaGrxB identified two iron-regulatory transcription factors, DaHapX and DaSreA, as interacting partners (Table S2; Figure S5, S6). These interactions were validated by Y2H and Co-IP (Fig 3A-C).

**Fig 3.**
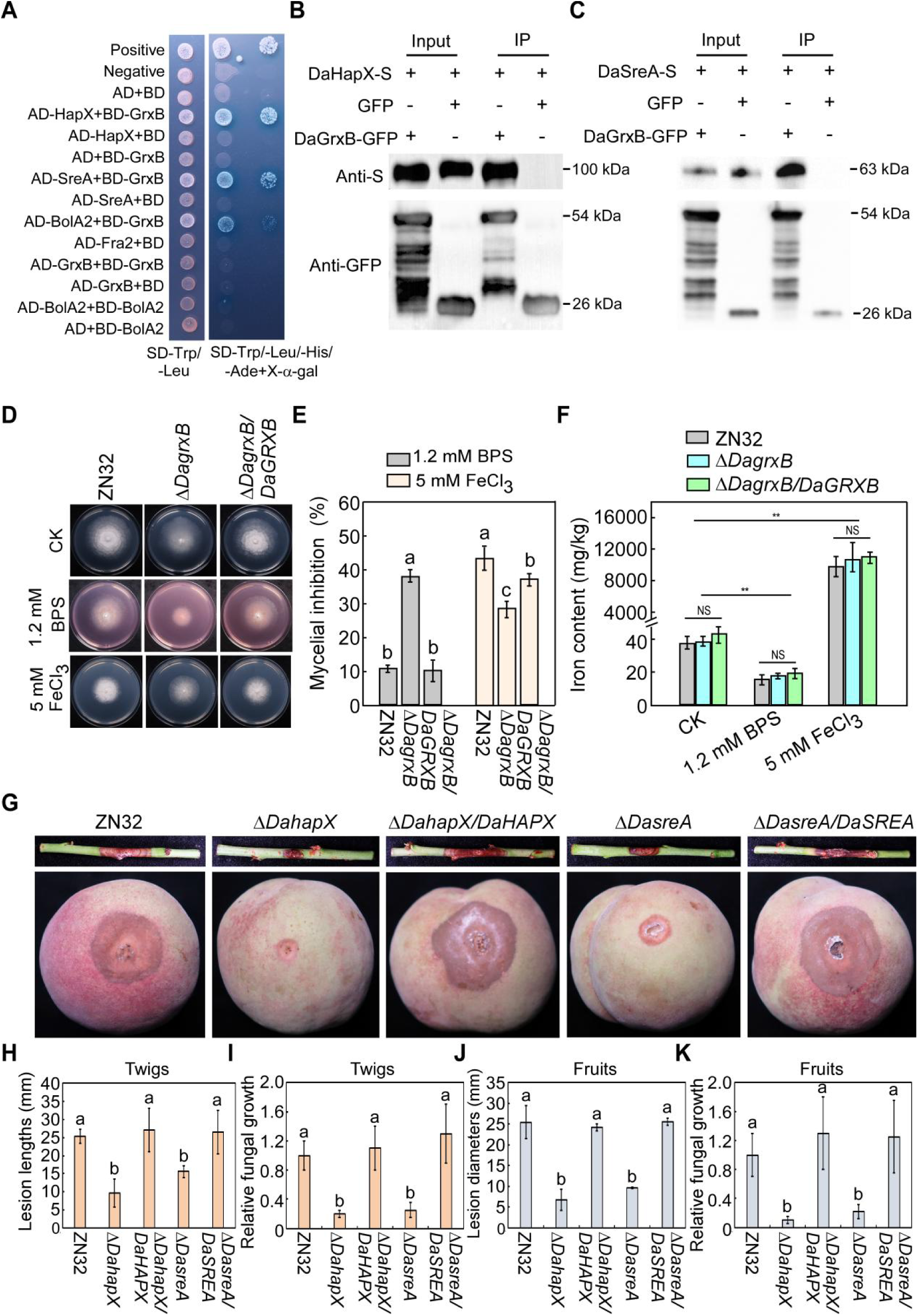
DaGrxB interacts with DaHapX and DaSreA and is required for iron stress responses. (A) Y2H assays. (B and C) Co-IP assays confirming interactions of DaGrxB with DaHapX (B) and DaSreA (C); GFP alone served as negative control. (D) Growth of ZN32, Δ*DagrxB* and complemented strain on CM with BPS (1.2 mM) or FeCl₃ (5 mM) for 4 d. (E) Mycelial inhibition rates. (F) Iron contents measured by AAS with or without iron stresses. NS, not significant; \*\**P* < 0.01. (G) Virulence assays of ZN32, Δ*DahapX*, Δ*DasreA* and complemented strains on twigs (7 d) and fruits (5 d). (H and J) Lesion lengths (H) and diameters (J). (I and K) Fungal biomass in twigs (I) and fruits (K) as in Figure 1C. Error bars, ±SD (n = 3). Different lowercase letters indicate significant differences (one-way ANOVA with Tukey’s HSD test, *P* < 0.05).

We next examined the growth of ZN32, Δ*DagrxB* mutants, and complemented strains under iron deficiency (1.2 mM BPS) and iron excess (5 mM FeCl₃). Δ*DagrxB* mutants exhibited more sensitive under BPS treatment, but showed enhanced tolerance to FeCl₃-mediated iron overload compared with the wild type and complemented strains (Fig 3D, E). To determine the specificity of this phenotype, we tested all four *GRX* deletion mutants under iron stresses. Among all four deletion mutants, only Δ*DagrxB* and Δ*DagrxD* showed increased sensitivity to BPS (∼40% and ∼20% inhibition, respectively), whereas only Δ*DagrxB* exhibited enhanced tolerance to iron excess (Fig S7A, B). Total cellular iron content did not differ between Δ*DagrxB* and ZN32 under any condition (Fig 3F).

To further establish the biological relevance of the DaGrxB-DaHapX/DaSreA interaction, we generated Δ*DahapX* and Δ*DasreA* deletion mutants (Fig S8A, B) and their complemented strains. Deletion of either *DaHAPX* or *DaSREA* significantly reduced virulence on both peach twigs and fruits (Fig 3G-K). These results demonstrate that DaGrxB, distinct from other family members, modulates iron homeostasis in *D. amygdali* through functional coordination with DaHapX and DaSreA

### DaGrxB, DaHapX, and DaSreA exhibit distinct localization patterns under different iron conditions

Under iron deficiency, Δ*DahapX* and Δ*DasreA* both showed increased sensitivity to 2.4 mM BPS, but only Δ*DahapX* exhibited altered colony morphology at 1.2 mM BPS. Δ*DagrxB* was sensitive to both 1.2 and 2.4 mM BPS. Under iron excess, Δ*DahapX* displayed enhanced resistance to 5 mM FeCl₃, similar to Δ*DagrxB*, while Δ*DasreA* showed increased sensitivity (Fig 4A).

**Fig 4.**
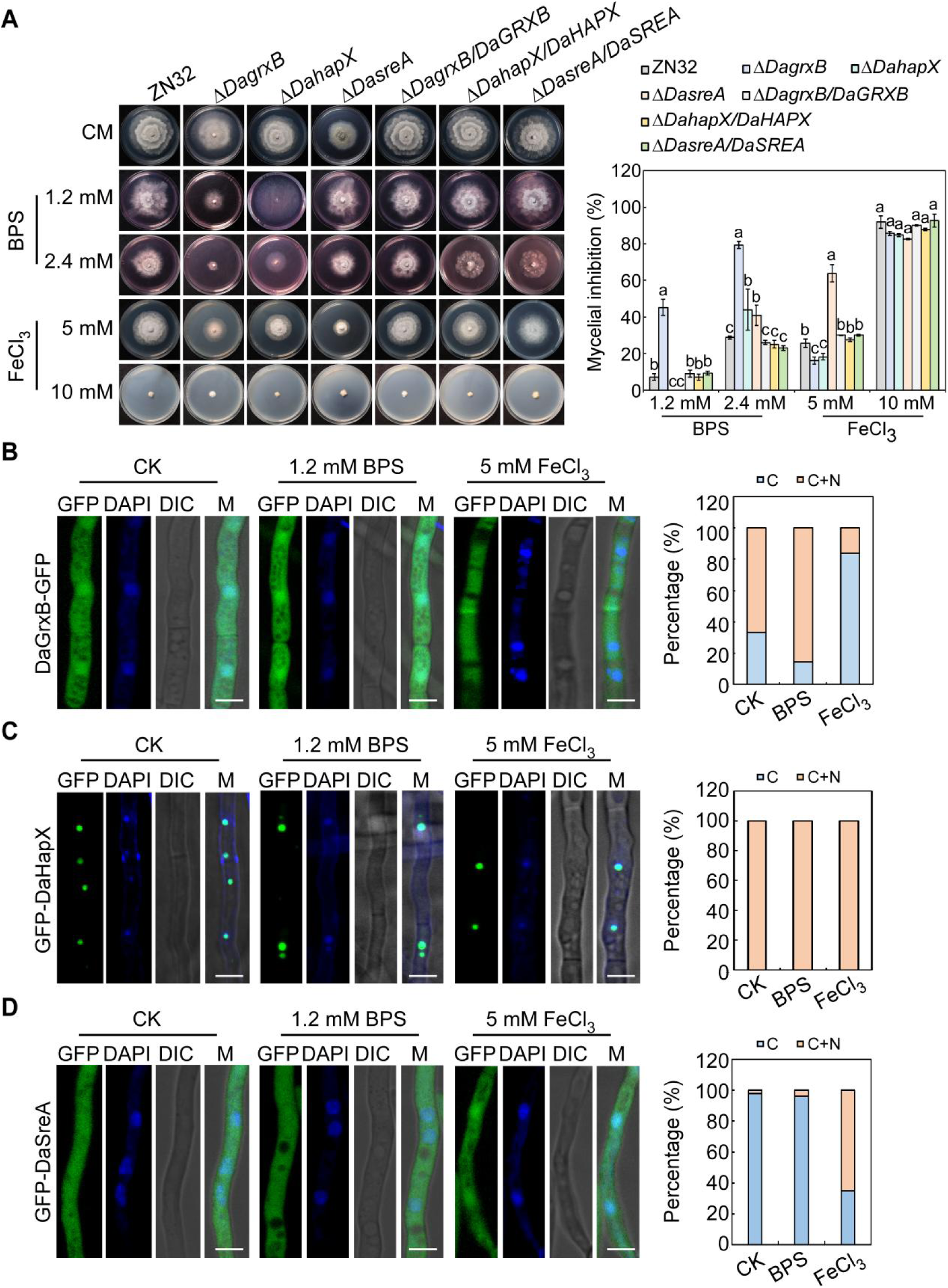
DaGrxB, DaHapX and DaSreA exhibit distinct localization patterns under different iron conditions. (A) Growth of ZN32, Δ*DagrxB*, Δ*DahapX*, Δ*DasreA* and complemented strains on CM with BPS (1.2, 2.4 mM) or FeCl₃ (5, 10 mM) for 4 d; untreated CM served as control. Bar graphs show mycelial inhibition rates. (B-D) Localization of GFP-DaGrxB (B), GFP-DaHapX (C) and GFP-DaSreA (D) under different iron conditions for 18 h; CK was under sufficient iron condition. DAPI was used as a nuclear marker. Percentages of cells with cytoplasmic versus nucleocytoplasmic localization are quantified in adjacent panels. Bars = 5 μm.

Under iron-sufficient conditions (CM media, CK), DaGrxB-GFP was distributed in both cytoplasm and nucleus in ∼67% of hyphae; under iron deficiency (1.2 mM BPS), this increased to 86% with enhanced nuclear fluorescence; under iron excess (5 mM FeCl₃), nuclear export was observed in 84% of hyphae (Fig 4B). DaHapX consistently localized to the nucleus regardless of iron status (Fig 4C). DaSreA was largely cytoplasmic under iron-sufficient and iron-deficient conditions, but partly entered the nucleus under iron excess (Fig 4D).

### DaGrxB, DaHapX, and DaSreA regulate iron homeostasis gene transcription under iron stresses

The expression of iron homeostasis-related genes was quantified in ZN32, Δ*DagrxB*, Δ*DahapX*, and Δ*DasreA* mutants under iron sufficiency (CM media, CK), iron deficiency (2.4 mM BPS), and iron excess (5 mM FeCl₃). Under iron-sufficient conditions, no significant differences in transcript levels were observed among the four strains (Fig 5A-M). Under iron deficiency, *DaHAPX* and five iron acquisition genes (*DaFREB*, *DaFETC*, *DaSIDA*, *DaSIDC*, *DaSIDD*) were strongly induced in ZN32; this induction was attenuated in Δ*DagrxB* and Δ*DahapX*, but unaffected in Δ*DasreA* (Fig 5A-H). Under iron excess, *DaSREA*, *DaCCCA*, and all tested iron utilization genes were induced in ZN32 (Fig 5C, I-M). *DaCCCA* induction was repressed in Δ*DagrxB* and Δ*DahapX* but retained in Δ*DasreA* (Fig 5I). DaGrxB, and DaHapX collectively contributed to the activation of iron utilization genes *DaACO1*, *DaCYCA*, *DaLYSF*, with the exception of *DaNAR1*(Fig 5J-M). DaSreA repressed iron acquisition genes, *DaHAPX* and *DaGRXB* under iron excess (Fig 5A, B, D-H). These results illustrate that under iron deficiency, DaGrxB and DaHapX coordinately activate iron acquisition while repressing *DaSREA*; under iron excess, DaHapX activates iron storage and utilization, whereas DaSreA represses iron acquisition as well as *DaHAPX* and *DaGRXB*.

**Fig 5.**
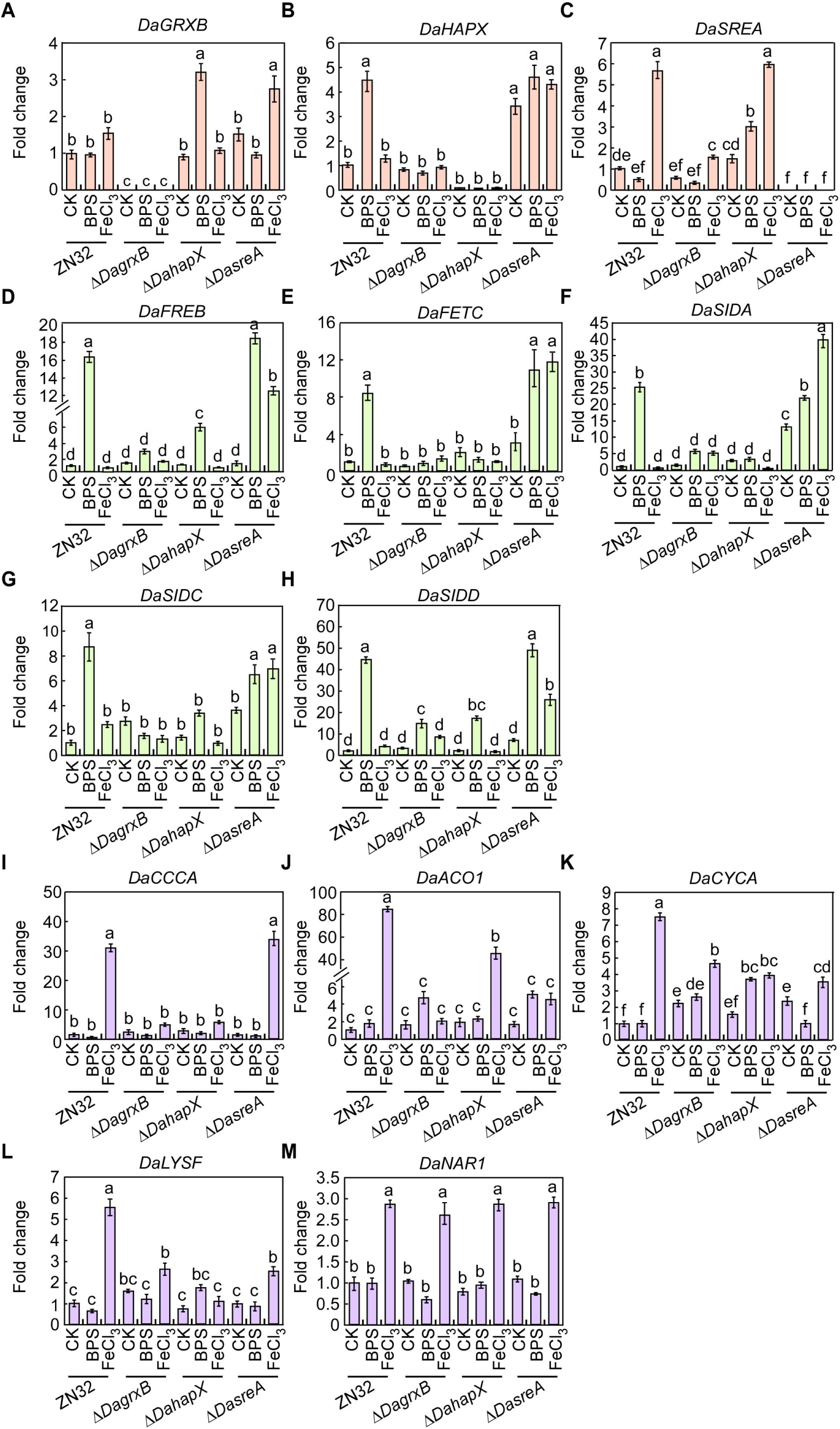
DaGrxB, DaHapX and DaSreA regulate iron homeostasis gene transcription under different iron conditions. Hyphal blocks of ZN32, Δ*DagrxB*, Δ*DahapX* and Δ*DasreA* were incubated in CM with or without iron stresses for 36 h. qRT-PCR was performed to detect expression of *DaGRXB* (A), *DaHAPX* (B), *DaSREA* (C), iron acquisition genes (*DaFREB* (D), *DaFETC* (E), *DaSIDA* (F), *DaSIDC* (G), *DaSIDD* (H), *DaCCCA* (I), and iron utilization genes (*DaACO1* (J), *DaCYCA* (K), *DaLYSF* (L), *DaNAR1*(M)). RNA from ZN32 without iron stress was used for normalization. Error bars, ±SD (n = 3). Different lowercase letters indicate significant differences (one-way ANOVA with Tukey’s HSD test, *P* < 0.05).

### *D. amygdali* dynamically coordinates iron homeostasis gene transcription across infection stages

To investigate iron dynamics during infection, we quantified total iron content in infected host tissues. Iron levels in infected peach fruits remained low at 1-2 days post-inoculation (dpi), increased significantly at 3-4 dpi, and then declined (Fig 6A). A similar trend was observed in twigs (Fig 6B). Notably, iron levels at 1-2 dpi were comparable to uninfected tissues (CK), indicating that the iron limitation during early infection is not host-induced deprivation, but an environment inherently insufficient for optimal pathogen growth. *DaGRXB* was strongly induced during early infection, peaking at ∼100-fold at 2 dpi; *DaHAPX* and *DaSREA* reached ∼25-fold and ∼12-fold (Fig 6C-E). Iron acquisition genes (*DaFTRA*, *DaFREB*, *DaFETC*, *DaMIRB*, *DaSIDA*, *DaSIDC*) and iron utilization genes (*DaACO1*, *DaERG11*, *DaHEMA*) were significantly upregulated at 1-2 dpi (Fig 6F-K, M, N). *DaCCCA* exhibited sustained upregulation from 3 to 7 dpi (Fig 6L), while *DaISA1* and *DaCYCA* were downregulated throughout infection (Fig 6P, Q). These results indicate that during early infection (1-2 dpi), *D. amygdali* encounters an iron-limited environment that triggers DaGrxB-dependent activation of DaHapX and subsequent induction of iron acquisition genes; by 3-4 dpi, host-mediated iron accumulation prompts activation of *DaCCCA* via DaHapX and repression of iron acquisition genes through DaSreA.

**Fig 6.**
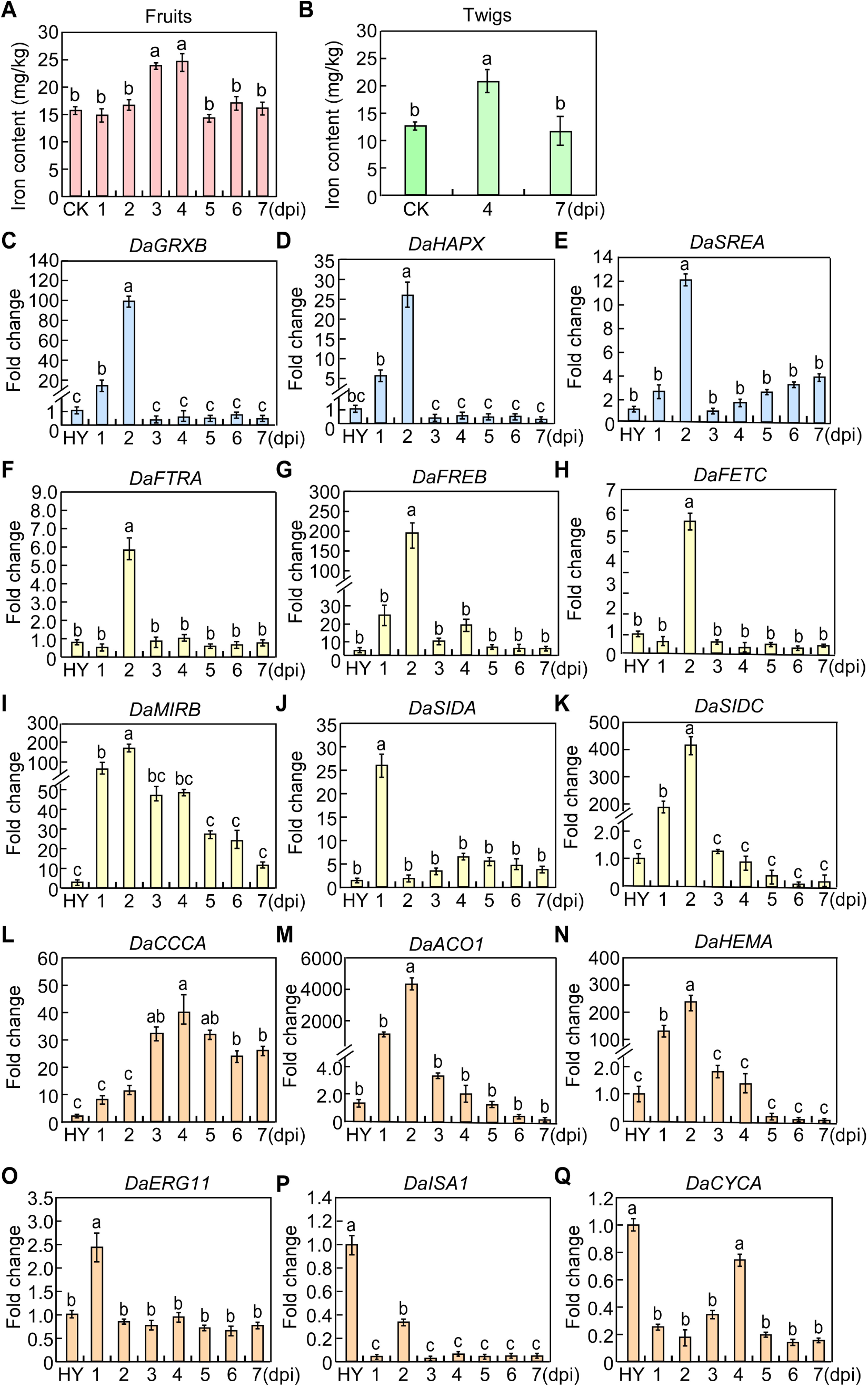
*D. amygdali* dynamically coordinates iron homeostasis gene transcription across infection stages. (A) Iron content in infected peach fruits at 1-7 dpi, measured by AAS. Uninfected fruits served as control (CK). (B) Iron content in infected peach twigs at 4 and 7 dpi. (C-Q) qRT-PCR analysis of *DaGRXB* (C), *DaHAPX* (D), *DaSREA* (E), iron acquisition genes (*DaFTRA* (F), *DaFREB* (G), *DaFETC* (H), *DaMIRB* (I), *DaSIDA* (J), *DaSIDC* (K), *DaCCCA* (L), and iron utilization genes (*DaACO1* (M), *DaHEMA* (N), *DaERG11* (O), *DaISA1*(P), *DaCYCA*(Q)) in infected fruits at 1-7 dpi. RNA from hyphae (HY) was used for normalization. Error bars, ±SD (n = 3). Different lowercase letters indicate significant differences (one-way ANOVA with Tukey’s HSD test, *P* < 0.05).

### Deletion of *DaGRXB* impairs Fe-S cluster biosynthesis and activates the antioxidant system

Cytosolic monothiol glutaredoxins serves as essential regulators and transfer agents for Fe-S clusters, delivering them to diverse Fe-S-dependent enzymes and proteins(Mühlenhoff et al., 2010; Berndt et al., 2021). To test whether DaGrxB contributes to Fe-S cluster biosynthesis, we quantified mitochondrial ferrous iron (Fe²⁺) levels using the Fe²⁺-specific fluorescent probe Mito-FerroGreen2. Δ*DagrxB* mutants exhibited significantly elevated mitochondrial Fe²⁺ accumulation (Fig 7A, B). Consistently, total, mitochondrial, and cytosolic aconitase (ACO) activities, a key [4Fe-4S] cluster-containing enzyme, were markedly diminished in Δ*DagrxB* (Fig 7C-E), and cellular ATP production was significantly reduced (Fig 7F), reflecting impaired mitochondrial respiratory function and defective Fe-S cluster biosynthesis. Surprisingly, Δ*DagrxB* mutants exhibited enhanced resistance to H₂O₂, t-BOOH, and diamide (Fig 7G, H). This resistance was accompanied by elevated GSH/GSSG ratio (Fig 7I), ∼4.0-fold increase in GR activity (Fig 7J), upregulated GST activity (Fig 7K), and markedly suppressed intracellular H₂O₂ levels (Fig 7L). Pretreatment with GSH or catalase significantly reduced wild-type virulence (Fig 7M, N). These results demonstrate that deletion of *DaGRXB* disrupts Fe-S cluster biosynthesis, and activates the GSH-dependent antioxidant system.

**Fig 7.**
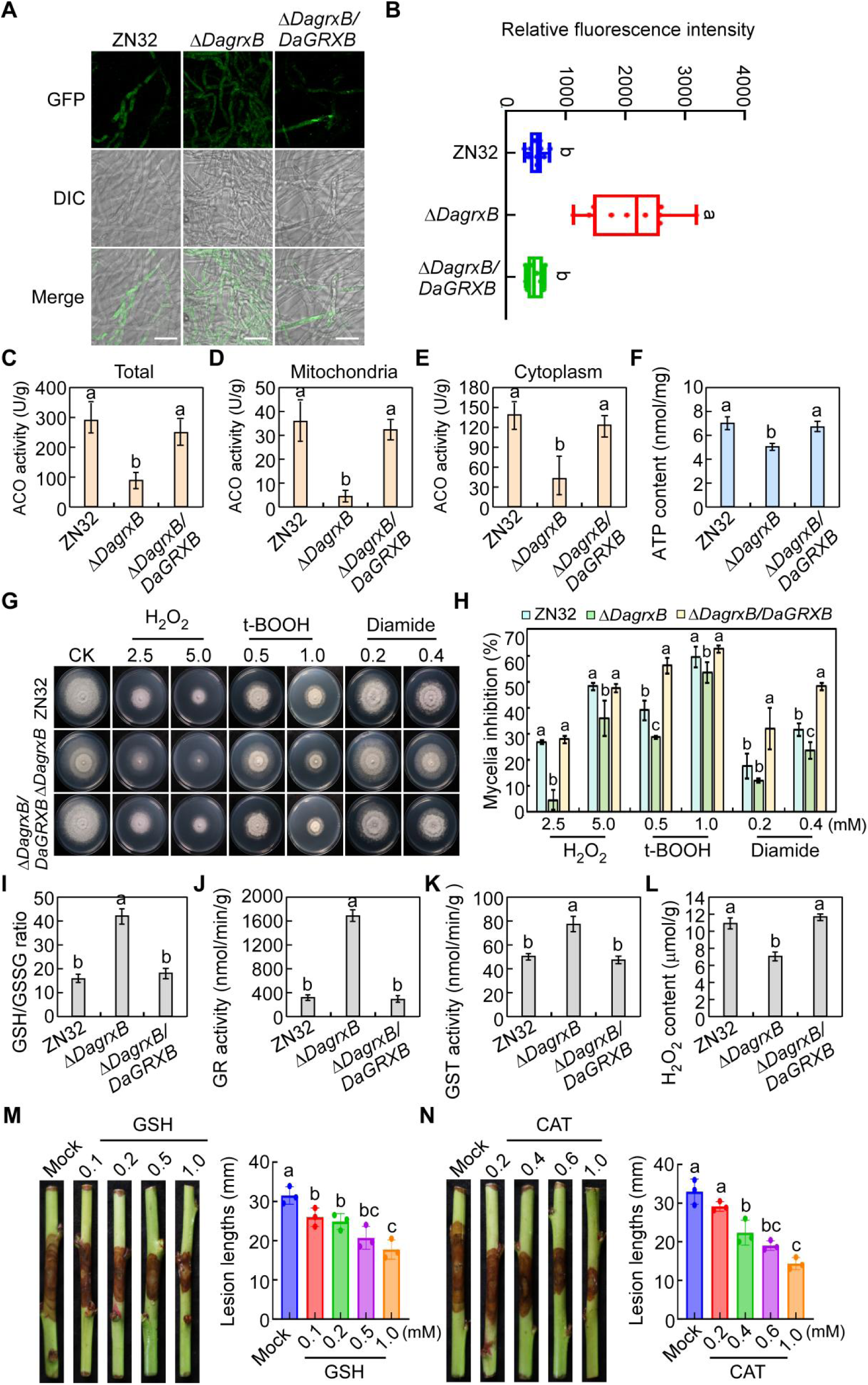
Deletion of *DaGRXB* impairs Fe-S cluster biosynthesis and activates the antioxidant system. (A) Confocal images of ZN32, Δ*DagrxB* and complemented strains stained with Mito-FerroGreen2 after 18 h in liquid CM. Bars = 20 μm. (B) Relative fluorescence intensity from (A) (n ≥ 50 images per strain). (C-E) Total, cytoplasmic and mitochondrial aconitase (ACO) activities. (F) ATP content. (G) Growth on CM with H₂O₂ (2.5, 5.0 mM), t-BOOH (0.5, 2.0 mM) or diamide (0.2, 0.4 mM) for 4 d. (H) Mycelial inhibition rates. (I) GSH/GSSG ratio. (J) GR activity. (K) GST activity. (L) Intracellular H₂O₂ content. (M) Disease symptoms and lesion lengths at 7 dpi after application of GSH (0.1-1.0 mM) followed by inoculation hyphal block of the wild type. (N) Same as (M) but with catalase (0.2-1.0 mM). Error bars, ±SD (n = 3). Different lowercase letters indicate significant differences (one-way ANOVA with Tukey’s HSD test, *P* < 0.05).

### The Trx domain cooperates with the Grx domain to mediate iron-responsive nucleocytoplasmic shuttling

Domain-specific deletion stains, Δ*DagrxB*/*DaGRXB*^ΔTrx^ (lacking the Trx domain) and Δ*DagrxB*/*DaGRXB*^ΔGrx^ (lacking the Grx domain), were generated to dissect the function of each domain. Phenotypic analyses demonstrated that both domain deletions recapitulated the full Δ*DagrxB* mutant phenotype, including abnormal colony growth, reduced pycnidia formation, and attenuated virulence (Fig 8A-D; S9A-E). Y2H confirmed intramolecular interaction between the Trx and Grx domains (Fig 8E). Both DaGrxB^ΔTrx^ and DaGrxB^ΔGrx^ exhibited diffuse cytoplasmic GFP fluorescence under iron sufficiency (CK). DaGrxB^ΔTrx^ retained nuclear localization under iron excess but remained cytoplasmic under iron deficiency (Fig 8F), whereas DaGrxB^ΔGrx^ showed constitutive cytoplasmic localization regardless of iron status (Fig 8G). These results indicate that Trx domain assists Grx domain in nucleocytoplasmic translocation through intramolecular interactions under different iron conditions, jointly regulated the development and virulence of *D. amygdali*.

**Fig 8.**
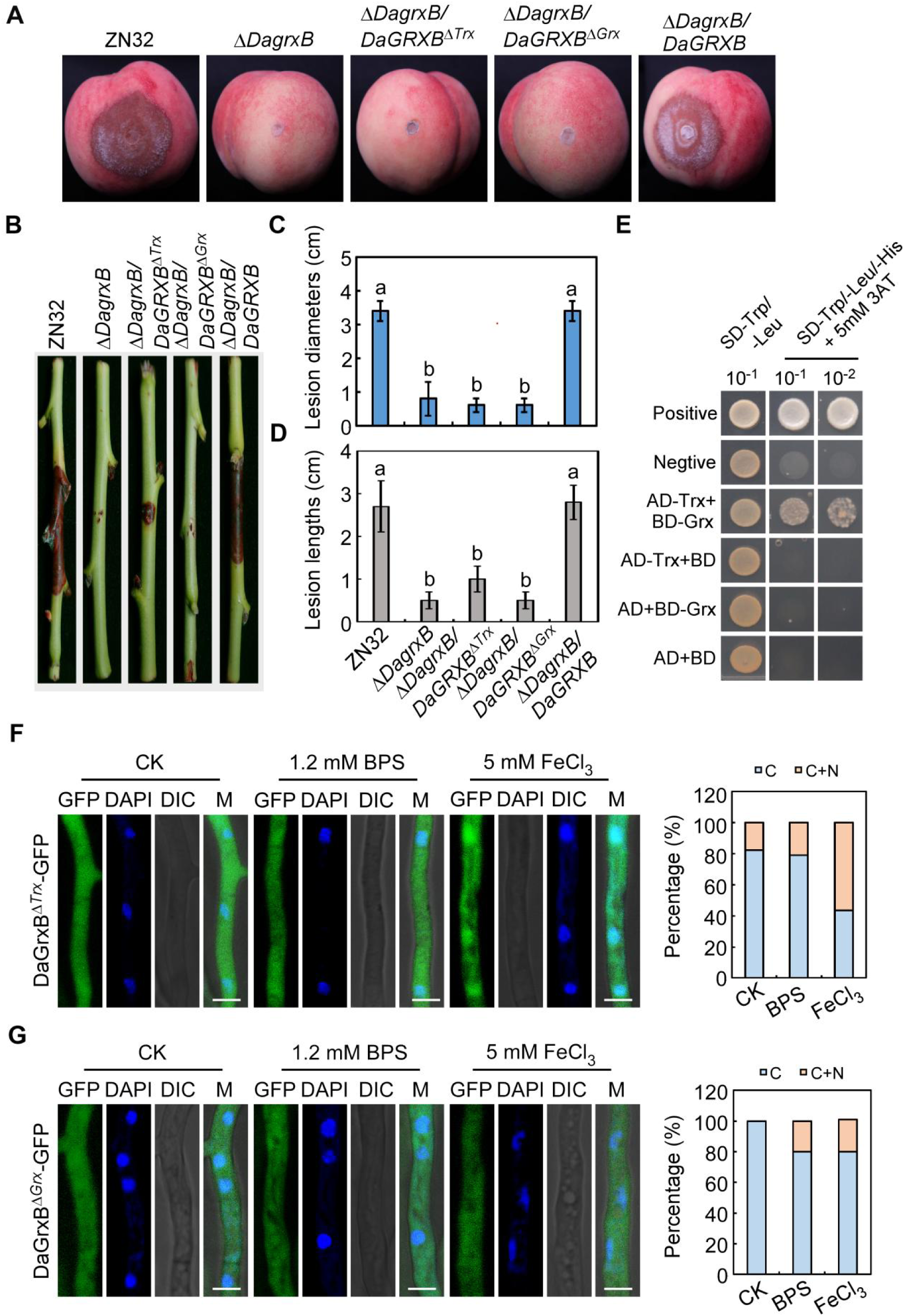
The Trx domain assists the Grx domain in nucleocytoplasmic translocation and virulence regulation. (A, B) Hyphal blocks of ZN32, Δ*DagrxB*, Δ*DagrxB*/*DaGRXB*^ΔTrx^, Δ*DagrxB*/DaGrxB^ΔGrx^, and complemented strains inoculated onto wounded peach fruits (A) or twigs (B). (C) Lesion diameters on fruits at 5 dpi. (D) Lesion lengths on twigs at 7 dpi. (E) Y2H assay confirming interaction between Trx and Grx domains. (F, G) Subcellular localization of DaGrxB^ΔTrx^-GFP (F) and DaGrxB^ΔGrx^-GFP (G) under different iron conditions; DAPI as nuclear marker. Percentages of cells with cytoplasmic versus nucleocytoplasmic localization quantified in adjacent panels. Bars = 5 μm. Error bars, ±SD (n = 3). Different lowercase letters indicate significant differences (one-way ANOVA with Tukey’s HSD test, *P* < 0.05).

### LDH-delivered dsRNA targeting *DaGRXB* suppresses *D. amygdali* infection

To establish a spray-induced gene silencing (SIGS) system against PSB, we synthesized dsRNA targeting *DaGRXB* (dsRNA*^DaGRXB^*), which significantly reduced disease symptoms (Fig S10A-D). The dsRNA*^DaGRXB^* was susceptible to RNase A but resistant to temperature and UV (Fig S11A-C). To improve efficacy, we complexed dsRNA*^DaGRXB^* with LDH at a 1:4 ratio, using dsRNA^GFP^-LDH as the negative control (Fig S12). Topical application of dsRNA*^DaGRXB^* alone to wounded peach twigs markedly attenuated PSB symptoms following *D. amygdali* inoculation, and dsRNA*^DaGRXB^*-LDH provided even stronger protection, as shown by reduced lesion development (Fig 9A, B) and fungal biomass (Fig 9C). Correspondingly, *DaGRXB* silencing ranged from 25-56% with dsRNA alone and from 31-91% with dsRNA*^DaGRXB^*-LDH (Fig 9D), confirming that LDH delivery enhanced RNAi efficiency. These findings suggest that LDH-mediated dsRNA delivery offers a promising strategy for PSB management.

**Fig 9.**
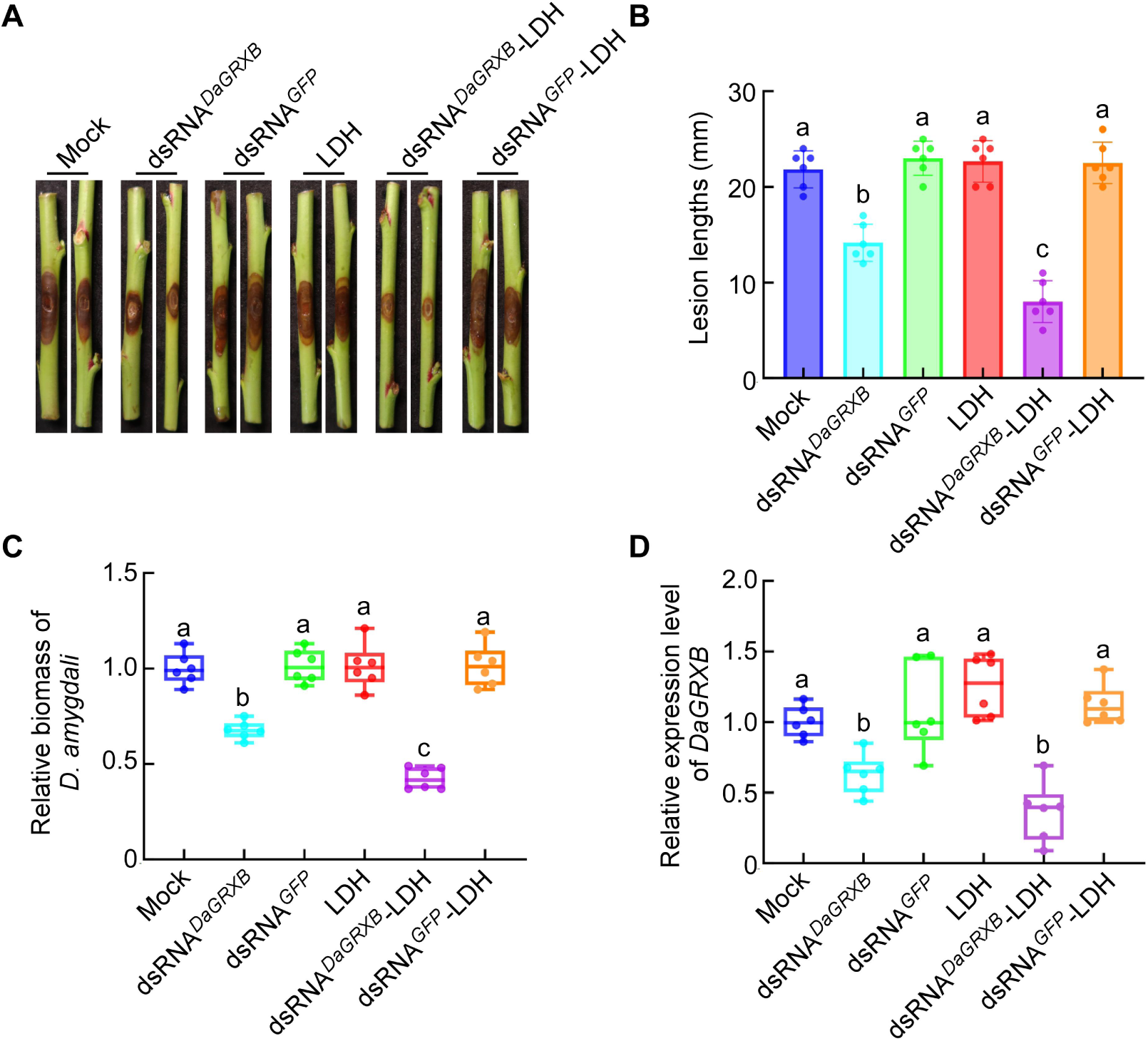
LDH-delivered dsRNA targeting *DaGRXB* suppresses *D. amygdali* infection. (A) Disease symptoms on peach twigs at 7 dpi after pre-treatment with sterile water (CK), dsRNA^DaGRXB^, dsRNA^GFP^, LDH, dsRNA^DaGRXB^-LDH, or dsRNA^GFP^-LDH for 2 h. (B) Lesion lengths. (C) Fungal biomass (qRT-PCR of *DaACTIN*, normalized to *PpTEF2*). (D) Relative expression of *DaGRXB* in infected tissues, normalized to *DaACTIN*. Error bars, ±SD (n = 3). Different lowercase letters indicate significant differences (one-way ANOVA with Tukey’s HSD test, *P* < 0.05).

## Discussion

We systematically characterized the function of four Grx proteins in *D. amygdali*. DaGrxB is the sole member essential for vegetative growth, asexual reproduction, and virulence. It interacts with iron-regulatory transcription factors DaHapX and DaSreA, orchestrating iron acquisition and storage via iron-dependent nucleocytoplasmic shuttling during infection. Deletion of *DaGRXB* disrupts Fe-S cluster biosynthesis and activates the GSH-dependent antioxidant system. Both Trx and Grx domains are indispensable for its function. Furthermore, LDH-delivered dsRNA targeting *DaGRXB* provides an effective RNAi-based strategy for controlling PSB.

The four Grx proteins show distinct subcellular localizations: DaGrxA and DaGrxC are mitochondrial, DaGrxB is nucleocytoplasmic, and DaGrxD shows stage-specific septal/cytoplasmic localization, broadly mirroring *S. cerevisiae*(Rodríguez-Manzaneque et al., 2002; Molina et al., 2004; Porras et al., 2006). Moreover, DaGrxA contains a typical CPYC motif characteristic of Class I dithiol Grx. DaGrxD possesses a CQFS motif homologous to the CPYS motif of found in certain dithiol Grxs, suggesting that they may retain deglutathionylating activity(Herrero et al., 2006). DaGrxB and DaGrxC harbor CGFS motif typical of Class II monothiol Grxs. Although all four *DaGRX* genes were upregulated by H₂O₂ treatment (Fig S13A-D), single-gene deletion of any *DaGRX* did not affect global glutathionylation level (Fig S13E), indicating functional redundancy among Grx family members in catalyzing reversible glutathionylation. However, the role of glutathionylation during infection remains to be explored in plant pathogenic fungi.

DaGrxB acts as an upstream regulator of DaHapX and DaSreA, physically interacting with both as well as with BolA2 (Fig S14), a cytoplastic Fe-S clusters biosynthesis regulator identified by MS. We propose the following model (Fig 10A): during early infection (1-2 dpi), iron in peach tissues is insufficient for optimal *D. amygdali* growth. DaGrxB accumulates in the nucleus, interacting with DaHapX to activate iron acquisition genes while repressing *DaSREA*; DaSreA remains cytoplasmic. At 3-4 dpi, host-mediated iron accumulation creates iron excess, prompting DaGrxB to exit the nucleus and interact with DaBolA2 to promote Fe-S cluster assembly. These clusters are transferred to DaHapX (activate iron storage) and to DaSreA (facilitating its nuclear entry, where it represses iron acquisition as well as *DaHAPX* and *DaGRXB*), forming a negative feedback loop. DaHapX and DaSreA are likely Fe-S proteins whose conformational changes respond to iron status, consistent with the findings in *A. fumigatus*(Lill et al., 2019).

**Fig 10.**
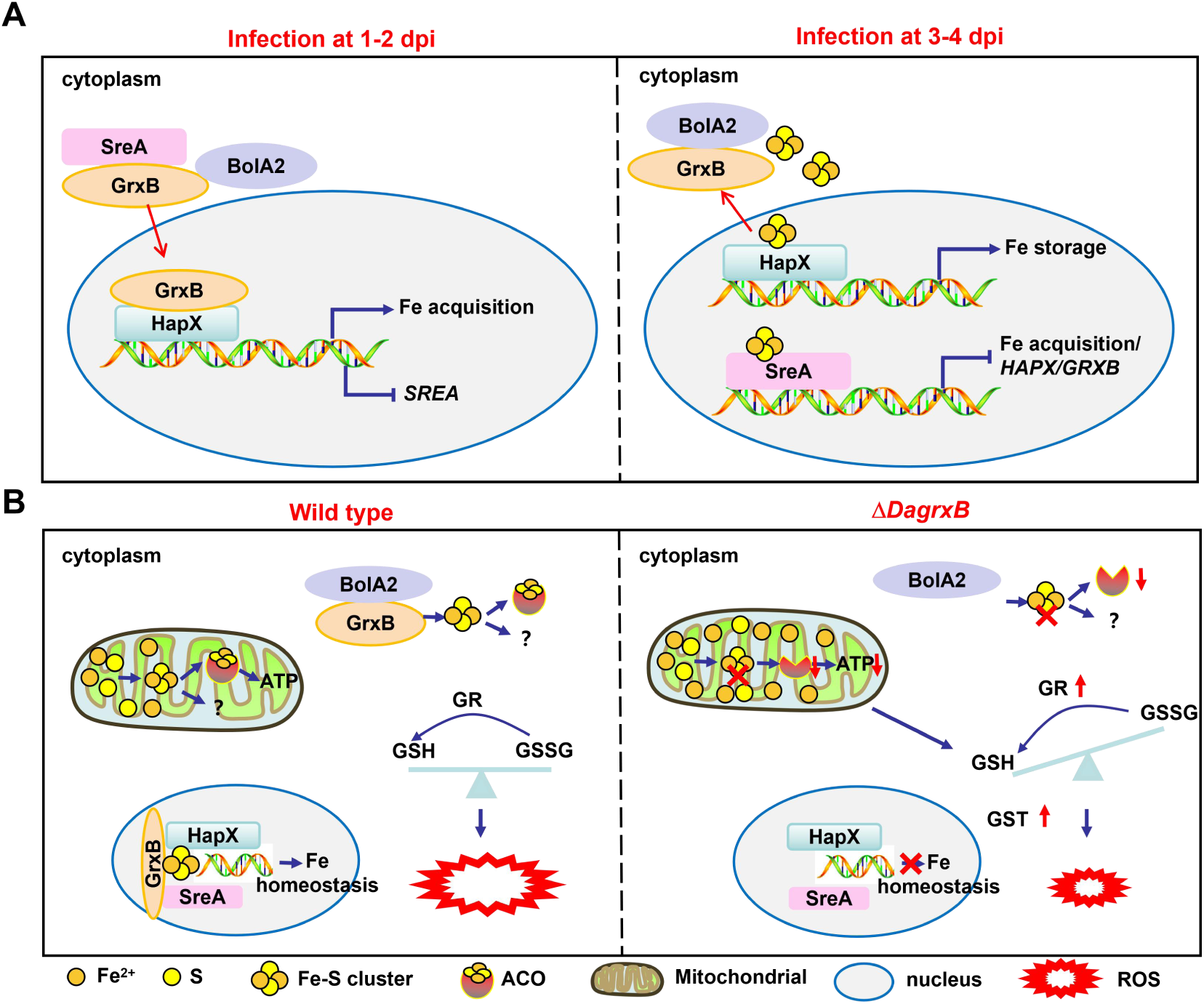
Working model of DaGrxB-mediated iron response and redox balance during *D. amygdali* infection. (A) During early infection (1-2 dpi), iron in peach tissues is insufficient for optimal growth. DaGrxB enters the nucleus to activate DaHapX-driven iron acquisition while repressing *DaSREA*; DaSreA remains cytoplasmic. At 3-4 dpi, host-mediated iron accumulation creates iron excess. DaGrxB exits the nucleus and interacts with DaBolA2 to assemble Fe-S clusters, which are transferred to DaHapX (activating iron storage) and DaSreA (triggering repression of iron acquisition genes as well as *DaHAPX* and *DaGRXB*), forming a negative feedback loop. (B) Deletion of *DaGRXB* leads to mitochondrial Fe²⁺ accumulation, impairing Fe-S cluster biosynthesis. Fe-S clusters are essential cofactors for numerous proteins, including ACO in the TCA cycle, respiratory chain complexes, and enzymes for DNA repair and amino acid biosynthesis. Consistent with disrupted Fe-S cluster assembly, total, mitochondrial, and cytosolic ACO activities were reduced in Δ*DagrxB*, accompanied by decreased ATP. Accumulated Fe²⁺ can react with H₂O₂ via the Fenton reaction to generate hydroxyl radicals. To prevent oxidative damage, Δ*DagrxB* constitutively activates the GSH-dependent antioxidant system, elevated GSH/GSSG ratios, increased GR and GST activities, and reduced H₂O₂, a compensatory response that protects against exogenous oxidants but imposes a metabolic cost that compromises virulence.

Several lines of evidence support DaGrxB as the primary iron sensor upstream of DaHapX and DaSreA. First, Δ*DagrxB* mutants are more sensitive to iron deficiency than Δ*DahapX* or Δ*DasreA*. Second, *DaGRXB* transcript levels are higher during early infection, and protein-level control appears to dominate over transcriptional changes. Temporal expression patterns indicate that *D. amygdali* transitions from an iron acquisition phase (1-2 dpi) to a phase marked by *DaCCCA* upregulation and repression of iron acquisition genes (3-4 dpi), coinciding with host-mediated iron accumulation. Similarly, Sun et al. (2023) reported low iron content in wheat heads at 1-2 dpi, increasing at 3-4 dpi following *F. graminearum* inoculation, accompanied by FgHapX-dependent *FgCCCA* activation and FgSreA-mediated repression of iron acquisition. Our results extend these findings, indicating that the pathogen might experience iron deficiency stress during the early infection phase.

The iron-dependent nucleocytoplasmic shuttling of DaGrxB resembles that of Grx4 in *C. neoformans*(Attarian et al., 2018). However, DaHapX is constitutively nuclear, whereas its homolog Hap43 in *C. albicans* enters the nucleus under iron deficiency and is exported under iron repletion(Snyder et al., 2020). DaSreA localizes to the cytoplasm under iron deficiency and enters the nucleus under iron excess, in contrast to Cir1 in *C. neoformans*, which remains nuclear under all conditions(Attarian et al., 2018). These differences highlight non-conserved dynamics, reflecting distinct evolutionary strategies for integrating iron sensing with transcriptional regulation. Given that deletion of either *DaHAPX* or *DaSREA* attenuates virulence, and iron acquisition genes are essential for virulence in the plant fungal pathogens(Eichhorn et al., 2006; López-Berges et al., 2012), we propose that impaired iron homeostasis is the primary cause of reduced virulence in Δ*DagrxB* mutants.

Deletion of *DaGRXB* impairs Fe-S cluster biosynthesis, aligning with the known function of monothiol glutaredoxins(Mühlenhoff et al., 2010; Berndt et al., 2021), and representing the first such report in a plant pathogenic fungus. Despite increased mitochondrial Fe²⁺ accumulation, Δ*DagrxB* mutants exhibited enhanced oxidative stress resistance, explained by the constitutive activation of the GSH-dependent antioxidant system, elevated GSH/GSSG ratios, upregulated GR and GST activities, and reduced intracellular H₂O₂ levels. Pretreatment with GSH or H₂O₂ removal suppressed pathogen virulence. We propose that mitochondrial Fe²⁺ accumulation triggers a compensatory antioxidant upregulation, which protects against exogenous oxidants but imposes a metabolic cost that compromises virulence. These findings suggest that the reduced virulence of Δ*DagrxB* mutants is also attributable to constitutively high GSH levels and low H₂O₂ levels, which may disrupt redox signaling required for successful host colonization (Fig 10B).

DaGrxB’s unique Trx domain accounts for its specialized function. Both domains are essential, and the Trx domain is required for the Grx domain’s iron-responsive shuttling via intramolecular interaction. In *S. cerevisiae*, both domains of ScGrx4 are indispensable for cell growth and Aft2 function, with the Trx domain enhancing GST activity of the Grx domain(Hoffmann et al., 2011; Chi et al., 2018). However, the Grx domain alone of GrxD is sufficient for iron-responsive nuclear shuttling in *A. fumigatus*(Lill et al., 2019), highlighting species-specific evolution of monothiol Grxs function.

RNAi-base disease management offers target specificity, environmental compatibility, and seamless integration with existing practices. RNAi targeting key virulence genes has been successfully employed, including *DCL1/DCL2* in *B. cinerea* (Wang et al., 2016), *ACTIN*, *VPS51*, and *SAC1* in *Rhizoctonia solani* and *Aspergillus niger*(Qiao et al., 2021), and *PMK1* in *M. oryzae* (Chen et al., 2025), as well as virulence effectors(Lee et al., 2024; Ouyang et al., 2024). *DaGRXB* is a promising RNAi target, as its deletion severely reduces virulence through critical roles in iron homeostasis and redox balance regulation. dsRNA targeting *DaGRXB* significantly lowers transcript levels and infection, with enhanced efficacy upon LDH complexation, consistent with advances in nanomaterial-assisted dsRNA delivery(Chen et al., 2025; Jiang et al., 2025; Guan et al., 2026).

Beyond application, the uncovered regulatory network may be conserved across plant pathogenic fungi, suggesting that targeting upstream iron homeostasis regulators could offer a broader disease-control strategy. Future work should focus on field applicability of this SIGS platform, including formulation stability, off-target effects, integration with existing practices, and efficacy under diverse environmental conditions. Understanding species-specific variations in iron regulation will be essential for developing tailored RNAi-based interventions against fungal pathogens.

## Supporting Information Legends

**Fig S1. Phylogenetic analysis, multiple sequence alignment, and domain architecture of fungal glutaredoxins.** (A) Phylogenetic tree constructed using the neighbor-joining (NJ) method with 1,000 bootstrap replicates in MEGA 7. Multiple sequence alignment visualized using ESPript 3.2. (B) Domain architecture of DaGrxs predicted by SMART.

**Fig S2. Transcriptional profiling of *DaGRX* genes across developmental and infection stages.** (A-D) Expression levels of *DaGRXA*-*DaGRXD* in vegetative hyphae (HY), pycnidia (PY), conidia (Co), and infected peach twigs at 12-72 hpi. Error bars, ±SD (n = 3). Different lowercase letters indicate significant differences (one-way ANOVA with Tukey’s HSD test, *P* < 0.05).

**Fig S3. PCR verification of single-gene deletion mutants of *DaGRXA*-*DaGRXD*.** Genomic DNA of ZN32 served as positive control; nuclease-free water as negative control.

**Fig S4. Subcellular localization of DaGrxA-DaGrxD.** (A-B) Localization of GFP-tagged DaGrxA-DaGrxD in hyphae (A) and conidia (B). (C) Colocalization of DaGrxA and DaGrxC with MitoTracker in hyphae. (D) Colocalization of DaGrxB with DAPI in hyphae and conidia. Bars = 5 μm.

**Fig S5. Phylogenetic analysis of fungal HapX proteins.** Tree constructed using the NJ method with 1,000 bootstrap replicates in MEGA 7.

**Fig S6. Phylogenetic analysis of fungal SreA proteins.** Tree constructed using the NJ method with 1,000 bootstrap replicates in MEGA 7.

**Fig S7. Phenotypic responses of *GRX* deletion mutants to iron stresses.** (A) Colony growth of ZN32, Δ*DagrxA*, Δ*DagrxB*, Δ*DagrxC*, Δ*DagrxD*, and complemented strains on CM with 1.2 mM BPS or 5 mM FeCl₃ for 4 d; CM only served as control. (B) Mycelial inhibition rates under iron stress. Error bars, ±SD (n = 3). Different lowercase letters indicate significant differences (one-way ANOVA with Tukey’s HSD test, *P* < 0.05).

**Fig S8. PCR verification of *DaHAPX* and *DaSREA* deletion mutants.**

Genomic DNA of ZN32 served as positive control; nuclease-free water as negative control.

**Fig S9. The Trx and Grx domains of DaGrxB are both essential for vegetative growth and asexual reproduction.** (A) Colony morphology of ZN32, Δ*DagrxB*, Δ*DagrxB*/*DaGRXB*^ΔTrx^, Δ*DagrxB*/*DaGRXB*^ΔGrx^, and complemented strains on PDA at 25°C for 4 d. (B) Pycnidia (15 and 30 d) and conidial morphology (40×; black triangles indicate β-conidia). Bar = 20 μm. (C) Colony diameters at 4 d. (D, E) Pycnidia numbers at 15 and 30 d. Error bars, ±SD (n = 3). Different lowercase letters indicate significant differences (one-way ANOVA with Tukey’s HSD test, *P* < 0.05).

**Fig S10. dsRNA targeting *DaGRXB* suppresses *D. amygdali* infection.** (A) Disease symptoms at 7 dpi following pre-treatment with sterile water (CK), dsRNA*^DaGRXB^*, or dsRNA*^GFP^* for 2 h. (B) Lesion lengths at 7 dpi. (C) Fungal biomass (qRT-PCR of *DaACTIN*, normalized to *PpTEF2*). (D) Relative expression of *DaGRXB* in infected tissues, normalized to *DaACTIN*. Error bars, ±SD (n = 3). Different lowercase letters indicate significant differences (one-way ANOVA with Tukey’s HSD test, *P* < 0.05).

**Fig S11. Stability of dsRNA*^DaGRXB^* under various treatments.** dsRNA*^DaGRXB^* (50 ng/μL) was incubated at 28°C for 0-20 d (A), exposed to UV for 0-8 h (B), or treated with 0.2 μg/mL RNase A for 0-150 min (C), then analyzed by agarose gel electrophoresis.

**Fig S12. Loading efficiency of dsRNA*^DaGRXB^* onto LDH nanoparticles at different mass ratios.** dsRNA*^DaGRXB^* (50 ng/μL) was complexed with LDH at ratios of 1:2, 1:4, 1:8, 1:16, and 1:20 (dsRNA : LDH), and analyzed by agarose gel electrophoresis. dsRNA*^GFP^* complexed at the same ratios served as control.

**Fig S13. Glutathionylation analysis of *DaGRX* deletion mutants.** (A-D) Transcript levels of *DaGRXA*-*DaGRXD* in ZN32 following treatment with 2.5 or 5.0 mM H₂O₂ for 0.5-2 h. (E) Western blot of protein glutathionylation in ZN32 and Δ*DagrxA*, Δ*DagrxB*, Δ*DagrxC*, Δ*DagrxD* mutants, with or without 2.5 mM H₂O₂ for 0.5 h. Proteins detected using anti-glutathione monoclonal antibody (ViroGen, #101-A) and HRP-conjugated anti-mouse secondary antibody. Error bars, ±SD (n = 3). Different lowercase letters indicate significant differences (one-way ANOVA with Tukey’s HSD test, *P* < 0.05).

**Fig S14. Phylogenetic analysis of fungal BolA-type regulators of Fe-S cluster biosynthesis.** Tree constructed using the NJ method with 1,000 bootstrap replicates in MEGA 7.

**Table S1 Primers used in this study.**

**Table S2 Partial list of proteins identified by Co-IP-MS as potential DaGrxB interactors.**

## Acknowledgements

We thank Wuhan Punaise Testing Technology Co., Ltd. for technical assistance with iron content measurement.

## Author contributions

Conceptualization: Zhaolin Ji, Lina Yang, You Liang.

Data curation: Xun Liu, Lili Xiong, Lina Yang.

Formal analysis: Yue Zhang, Yaru Jing.

Funding acquisition: Zhaolin Ji, Lina Yang.

Investigation: Hengsong Shi, Ziyi Pan, Shiyuan Cao.

Methodology: Xun Liu, Lili Xiong, Lianwei Li, You Liang, Lina Yang.

Project administration: Zhaolin Ji, Lina Yang.

Resources: Zhaolin Ji. Software: Lina Yang.

Supervision: Zhaolin Ji, Lina Yang.

Validation: Xun Liu, Lili Xiong, Lina Yang.

Writing-original draft: Lina Yang.

Writing-review & editing: Lina Yang, Lianwei Li.

